# Reconstruction of ancestral protein sequences using autoregressive generative models

**DOI:** 10.1101/2024.09.17.613387

**Authors:** Matteo De Leonardis, Andrea Pagnani, Pierre Barrat-Charlaix

## Abstract

Ancestral sequence reconstruction (ASR) is an important tool to understand how protein structure and function changed over the course of evolution. It essentially relies on models of sequence evolution that can quantitatively describe changes in a sequence over time. Such models usually consider that sequence positions evolve independently from each other and neglect epistasis: the context-dependence of the effect of mutations. On the other hands, the last years have seen major developments in the field of generative protein models, which learn constraints associated with structure and function from large ensembles of evolutionarily related proteins. Here, we show that it is possible to extend a specific type of generative model to describe the evolution of sequences in time while taking epistasis into account. We apply the developed technique to the problem of Ancestral Sequence Reconstruction (ASR): given a protein family and its evolutionary tree, we try to infer the sequences of extinct ancestors. Using both simulations and data coming from experimental evolution we show that our method outperforms state-of-the-art ones. Moreover, it allows for sampling a greater diversity of potential ancestors, allowing for a less biased characterization of ancestral sequences.

## I. INTRODUCTION

Homologous proteins have a common evolutionary origin that can go back to billions of years. Throughout their evolution, they diversify through mutations while selection preserves their biological function. Consequently, many protein families contain thousands of sequences that are highly variable and yet maintain similar structures and functions. On the other hand, even a few mutations can destabilize a protein and destroy its function. A quantitative description how protein sequences change in time is thus a challenging problem, with important consequences for our understanding of the evolution of life.

Many probabilistic models of protein sequence evolution have been developed. Commonly used ones describe the evolution at each sequence position as a Markov chain across amino acid states, taking into account average properties of the substitution process such as more frequent transitions between similar amino acids [1–3]. Variations in evolutionary speed at different sites are often represented by using a set of substitution rates to which sites can be assigned, usually coming from a Gamma distribution [4]. An important and widely accepted assumption is that sequence positions evolve independently. This has the advantage of greatly simplifying sequence evolution models, making them convenient to manipulate analytically and computationally manageable. However, it comes at the cost of ignoring epistasis, that is the fact that the effect of a mutation depends on the rest of the sequence.

Sequence evolution models are used in the general field of phylogenetics which explores the evolutionary relations between proteins. An notable application is that of ancestral sequence reconstruction (ASR): given a set of homologous sequences and their phylogenetic tree, ASR consists in inferring likely sequences for the internal nodes of the tree, which correspond to extinct ancestral proteins. Reconstructed proteins can then be synthesized and tested in the lab. The technique is used to study the sequence-function relationship in proteins, for instance by understanding which mutations cause a change in enzymatic activity or binding specificity of a protein [5–7]. It can also be used to address fundamental evolutionary questions, such as the evolution reaction specificity or thermostability of proteins across the tree of life [8, 9].

The large amount of protein sequence data combined with recent theoretical and computational work has also allowed the development of generative protein sequence models. These models build on the idea that the sequence variability among homologous protein with similar biological functions inform us about the sequence-function relationship. In practice, generative models are trained using large amounts of protein sequences and consist of a probability distribution *P*(**s**) over any potential amino acid sequence, with functional ones presumably being more probable. Classes of models include ones inspired from statistical physics such as the Potts model [10] and restricted Boltzmann machines [11], or based on neural networks such as transformers [12, 13]. A major achievement of these models is the possibility of using them to sample new artificial sequences that are distant from any natural protein but still functional [14, 15].

An essential ingredient for the success of generative model is the modeling of *epistasis*: the fact that the effect of a mutation on protein function depends on the rest of the sequence. Epistasis is caused by interaction between amino acids, and is essential to describe the fitness landscape of a protein [16, 17]. Interestingly, it has also been suggested that epistasis may be the cause of variable evolutionary rates across phylogenetic trees [18]. Since common sequence evolution models ignore epistasis, they can only represent a crude approximation of the evolutionary constraints acting on a protein. As the change of a protein sequence in time depends on functional constraints, it is reasonable to expect that an inaccurate representation of the fitness landscape negatively affects the modeling of dynamics. On the other hand, the non-independence of mutations that characterizes generative models makes it challenging to use them for dynamical purposes, as most existing algorithms assume independence of positions. Different studies have proposed using the Potts model to describe evolutionary dynamics, but current techniques allow for little analytical treatment and are limited to forward simulation of sequences [19, 20]. Consequently, they cannot be used for tasks such as inference of phylogenies or ancestral sequence reconstruction.

In this study, we set out to extend the application of generative models to describe evolutionary dynamics. First, we develop an analytically and numerically tractable sequence evolution model with generative properties, based on the so-called ArDCA generative model and its autoregressive architecture [21]. Our model accounts for epistasis and is generative over long-term evolution, but also allows use of some of the standard algorithms used in phylogenetics such as Felsenstein’s pruning algorithm [22]. We then apply our model to ancestral sequence reconstruction (ASR) and demonstrate, using simulated data, that it outperforms the state-of-the-art program IQ-TREE [23]. Finally, we validate our approach with recent experimental data on directed evolution and show that reconstruction of a known ancestor is done more accurately than using IQ-TREE. To our knowledge, this is the first use of such data to evaluate reconstruction methods.

## II. RESULTS

### A. Autoregressive model of sequence evolution

Models of evolution commonly used in phylogenetics rely on the assumptions that sequence positions evolve independently and that evolution at each position *i* follows a continuous time Markov chain (CTMC) parametrized by a substitution rate matrix **Q**^*i*^. Matrix **Q**^*i*^ is of dimensions *q* × *q* where *q* = 4 for DNA, 20 for amino acids or 64 for codon models. The probability of observing a change from state *a* to state *b* during evolutionary time *t* is then given by 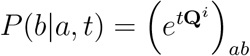.

If the model is time-reversible, it is a general property of CTMCs that the substitution rate matrix can be written as

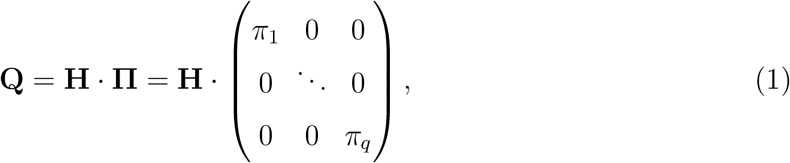

where **H** is symmetric and **Π** is diagonal with entries that sum to 1 [24]. The two matrices have simple interpretations. On the first hand, **Π** fixes the long-term equilibrium frequencies, that is 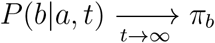. On the other, **H** influences the dynamics of the Markov chain but does not change the equilibrium distribution. Most commonly, both matrices are considered to be independent of the sequence position *i*, and **H** can potentially be scaled in order to represent different rates of evolutionary change [4].

In order to incorporate constraints coming from a protein’s structure and function into the evolutionary model, we develop a protein family specific model of protein sequence evolution based on the the autoregressive generative model ArDCA [21]. ArDCA models the diversity of sequences in a protein family using a set of learned conditional probabilities. In practice, the model assigns a probability to any sequence **a** = {*a*_1_, …, *a*_*L*_} of *L* amino acids:

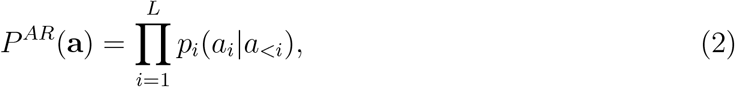

where the product runs over positions *i* in the sequence of length *L* and *a*_<*i*_ = *a*_1_, …, *a*_*i*−1_ represents the amino acid states before position *i*. Functions *p*_*i*_ represent the probability according to the model to observe state *a*_*i*_ in position *i*, given that the previous amino acids were *a*_1_, …, *a*_*i*−1_. They are learned using the aligned sequences of members of the family. Their precise functional form remains rather simple and is given in the methods section. Note that the autoregressive architecture is also employed in the context of deep-learning methods, to which the model we describe below could potentially be generalized [13]. In actual implementations, the order in which the product in Eq. 2 is performed is not the natural (1, …, *L*) but rather an order where positions are sorted by increasing variability. This does not significantly affect the model we present below, and we keep the notation of Eq. 2 for simplicity.

It has been shown in [21] that the generative capacities of ArDCA are comparable to that of state of the art models such as bmDCA [17]. This means that a set of sequences sampled from the probability in Eq. 2 is statistically hard to distinguish from the natural sequences used in training or, in other words, that the model can be used to sample new artificial homologs of a protein family. The generative capacities of a protein model comes from its ability to represent epistasis, that is the relation between the effect of a mutation and the sequence context in which it occurs. Here, epistasis is modeled through the conditional probabilities *p*_*i*_: the distribution of amino acids at position *i* depends on the states at the previous positions 1, … *i* − 1.

We take advantage of the autoregressive architecture to define a generative evolution model. Given two amino acid sequences **a** and **b**, we propose that the probability of **a** evolving into **b** in time *t* take the form

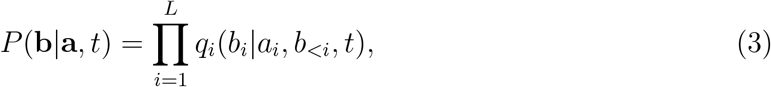

where the position specific conditional propagator *q*_*i*_ is defined as

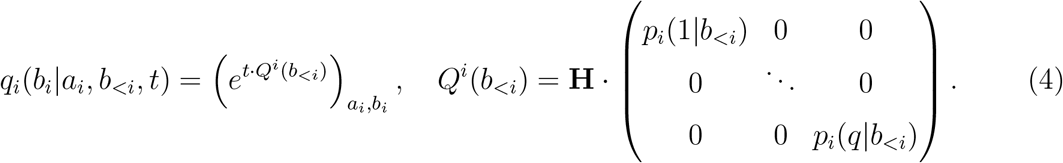

According to these equations, evolution for each position *i* follows a standard CTMC. However, we use the decomposition of Eq. 1 to set the equilibrium frequency at *i* to *p*_*i*_(*b*|*b*_<*i*_). In other words, we consider that position *i* evolves in the context of *b*_1_, …, *b*_*i*−1_, and that its dynamics are constrained by its long term frequency given by the autoregressive model. An important consequence of this choice is that our evolution model will converge at long times to the generative distribution *P*^*AR*^:

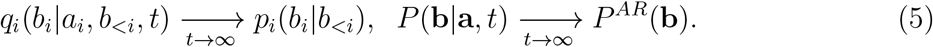

We argue here that such a property is essential to build a realistic protein sequence evolution model, particularly when considering evolution over long periods. Note that to converge to a generative distribution, accurate modeling of epistasis is required. Using site-specific frequencies would not be sufficient, as the effect of mutations in a protein sequence typically depends on the context [16]. The technique proposed here allows us to represent epistasis through the context dependent probabilities *p*_*i*_, while still considering each sequence position one at a time.

Interestingly, we note that the model in Eq. 3 is not time reversible, although context dependent site propagators in Eq. 4 are reversible. We show in the Supplementary Material that this is mainly an artifact of the autoregressive nature of the model coupled with epistasis. Using non-time reversible evolutionary models is uncommon in the field, but this is mainly due to practical considerations and there are no fundamental reasons for evolution itself to be reversible [22]. In practice, this means that algorithms using this model have to be adapted accordingly.

We underline that this approach has important differences with standard models of evolution used in phylogenenetics. In phylogenetic reconstruction, the tree and the sequence evolution model are usually inferred at the same time and from the same data. The number of parameters of the evolution model is then kept low to reduce the risk of overfitting, for instance by using a predetermined set of evolutionary rates to account for variable and conserved sites. Methods that introduce more complex models such as site specific frequencies do so by jointly inferring the parameters and the tree, leading to computationally intensive algorithms [25, 26].

Here instead, parameters of the generative model in Eq. 2 are learned from a protein family, *i*.*e*. a set of diverged homologous protein sequences. While it is true that these sequences share a common evolutionary history and cannot be considered as independent samples, common learning procedures only account for this in a very crude way [10, 21]. Despite this, it appears that the generative properties of such models are not strongly affected by the phylogeny [27, 28]. This allows us to proceed in two steps: first construct the model from data while ignoring phylogeny, and then use it for phylogenetic inference tasks.

An advantage of this approach is that once the model of Eq. 2 is inferred, the propagator in Eq. 3 comes “for free” as no additional parameters are required. Importantly, our model does not use site specific substitution rates. Indeed, it has been shown that these can be seen as emergent properties of more complex models of evolution [18]. However, a constraint is that the inference of the generative model requires the existence of an appropriate training set, that is a protein family with sufficient variability among its members.

### B. Ancestral sequence reconstruction

We apply our evolutionary model to the task of ancestral sequence reconstruction (ASR). The goal of ASR is the following: given a set of extant sequences with a shared evolutionary history and the corresponding phylogenetic tree, is it possible to reconstruct the sequences of extinct ancestors at the internal nodes of the tree? Along with the autoregressive evolutionary model described above, we thus need two inputs to perform ASR: a known phylogenetic tree, and the multiple sequence alignment of the leaf sequences. The length of the aligned sequences has to exactly correspond to that of the autoregressive model.

To reconstruct ancestral sequences using the autoregressive model, we proceed as follows:

i. for sequence position *i* = 1, use the evolution model defined by the equilibrium frequencies *p*_1_ to reconstruct a state 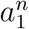 at each internal node *n* of the tree;
ii. iterating through subsequent positions *i* > 1: reconstruct state 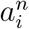 at each internal node *n* using the model defined in Eq. 4, with the context 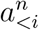 having been already reconstructed in the previous iterations.

It is important to note that when any position *i* > 1 is reconstructed, the context at different internal nodes of the tree may differ. For a branch joining two nodes (*n, m*) of the tree, the evolution model will thus differ if we go down or up the branch: in one case the context at node *n* must be used, in the other case the context at node *m*. This is a consequence of the time-irreversibility of the model. For this reason, we use a variant of Felsenstein’s pruning algorithm that is adapted to irreversible models [29]. This comes at no computational cost.

Note that this method is adapted to both maximum likelihood and Bayesian inference. In the ML case, each iteration reconstructs the most probable residue is at a position *i* given the already reconstructed context. In the Bayesian case, residue 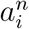 is instead sampled from the posterior distribution.

In any realistic application, the phylogenetic tree has to be reconstructed from the aligned sequences. In principle, a consistent approach would use the same evolutionary model for tree inference and ASR. However, our model does not allow us to reconstruct the tree. Therefore, in any realistic application, the tree is reconstructed using an evolutionary model that typically will differ from ours. To reduce issues related to this evolutionary model discrepancy, we adopt the following strategy: our ASR method blindly trusts the topology of the input tree, but recomputes the branch length using the sequences. As explained in the Methods, there is no direct way to optimize branch length with the autoregressive model. For simplicity, we use profile model with position-specific amino acid frequencies for this task. This provides a relatively accurate estimate of the branch lengths, as shown in Figure S2.

### C. Results on simulated data

There are two difficulties when evaluating the capacity of a model to perform ASR. The first is that in the case of biological data, the real phylogeny and ancestral sequences are usually not known. As a consequence, one must rely on simulated data to measure the quality of reconstruction. The second is that the reconstruction of an ancestral sequence is always uncertain, as evolutionary models are typically stochastic. The uncertainty becomes higher for nodes that are remote from the leaves. This means that it is only possible to make a statistical assessment about the quality of a reconstruction.

To test our approach, we adopt the following setup. We first generate phylogenetic trees by sampling from a coalescent process. We decide to use Yule’s coalescent instead of the more common Kingman. The latter tends to produce a large majority of internal nodes in close vicinity to the leaves with the others separated by very long branches, resulting in a trivial reconstruction for most nodes and a very hard one for the deep nodes. Yule’s coalescent generates a more even distribution of node depths, allowing us to better evaluate reconstruction quality, see Supplementary Material and Figure S3. For each tree, we simulate the evolution of sequences using a model that we refer to as “evolver” to obtain two multiple sequence alignments, one for the leaves and one for the internal nodes of the tree. We then reconstruct internal nodes using the desired approach by using the leaf alignment and the tree topology as input data.

We will consider two kinds of evolver models: *(i)* the same autoregressive model that we will then use for reconstruction, which is an ideal case and *(ii)* an evolutionary model based on a Metropolis sampling of a Potts model. These two evolvers come from models trained on actual protein families: we use evolvers based on the PF00072 response regulator family for results of the main text, and show results for three other families (PF00014, PF00076 and PF00595) in the Supplementary Material. It is important to note that the approach that we propose only makes sense when considering the evolution a protein family on which the model in Eq. 2 is trained. Hence, any evolver model used in our simulations should reproduce at long times the statistics of the considered protein family, *i*.*e*. it should satisfy Eq. 5. For this reason, we only consider the two evolvers above and do not use more traditional evolutionary models such as an arbitrary GTR on amino-acids [30].

For reconstruction, we compare our autoregressive approach to the commonly used IQ-TREE program [23]. When supplied with a protein sequence alignment and a tree, IQ-TREE infers a joint substitution rate matrix for all sequence positions, with rates that can differ across positions. Both methods run on a fixed tree topology, with branch lengths being re-inferred using maximum likelihood (see Methods).

#### Autoregressive evolver

We first investigate the case of the autoregressive evolver. This setting is of course ideal for our method, as there is perfect coincidence between the model used to generate the data and to perform ASR. We first evaluate the quality of reconstruction by computing the Hamming distance of the real and inferred sequences for each internal node of the simulated phylogenies. The left and central panels of Figure 1 show this Hamming distance as a function of the node depth, that is the distance separating the node from the leaves, and for a maximum likelihood reconstruction. Hamming distance is computed including gap characters in the aligned sequences on the right panel, while they are ignored on the central one. We see that the autoregressive reconstruction clearly outperforms the state of the art method: the improvement in Hamming distance increases with node depths, and the distance to the real ancestor drops from ∼ 0.4 to ∼ 0.3 when using the autoregressive approach. The increase in reconstruction quality with node depths is consistent with recent findings that epistasis only becomes important at relatively large sequence divergences [20, 31].

**FIG. 1.**
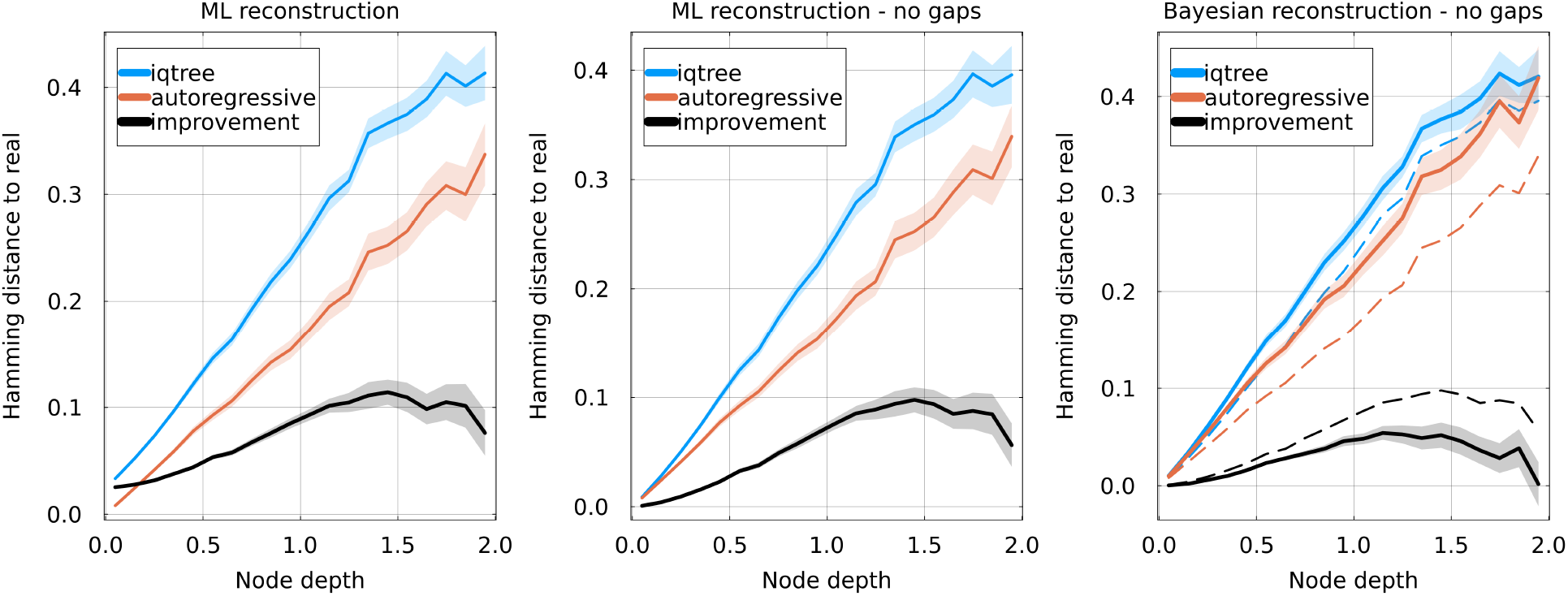
Hamming distance between reconstructed and real sequences as a function of node depth, using IQ-TREE and our autoregressive approach. The difference between the two methods (“improvement”) is shown as a black curve. Estimation of the incertitude is shown as a ribbon. The evolver and reconstruction autoregressive models are learned on the PF00072 family. **Left**: Hamming distance between the full aligned sequences, gaps included, using maximum likelihood reconstruction. **Center**: Hamming distance ignoring gapped positions, using maximum likelihood reconstruction. **Right**: comparison of Bayesian (solid lines) and maximum likelihood (dashed lines) reconstructions, ignoring gaps.

Interestingly, the performance of IQ-TREE degrades if Hamming distance is computed including gaps, as in the left panel. This is because like other popular methods, IQ-TREE treats gaps in input sequences as unknown amino acids, and reconstructs an ancestral amino acid for gapped positions [23, 32]. On the contrary, our autoregressive approach treats gaps as if they were an additional amino acid and will reconstruct ancestral sequences that can contain gaps. This benefits the autoregressive approach as aligned ancestral sequences can in fact contain gaps. This effect is particularly visible at low node depths. However, ignoring the effects of gaps in the Hamming distance also leads to a clear improvement when using the autoregressive approach as shown in the central panel.

The right panel of Figure 1 shows the quality of the reconstruction for Bayesian reconstruction. In this case, a ensemble of sequences is reconstructed for each internal node, and the metric is the average Hamming distance between this ensemble and the real ancestor. Gaps are again ignored when computing the Hamming distance. We again observe an improvement when using the autoregressive method, of slightly lesser magnitude than in the maximum likelihood case.

#### Properties of reconstructed sequences

To further analyze the reconstructed sequences, we first look at the diversity of generated ancestors in Bayesian reconstruction. The left panel of Figure 2 shows the average Hamming distance between sequences reconstructed at the same internal node, as a function of depth. For deeper nodes (depth ≳ 1), the Bayesian autoregressive approach reconstructs a significantly more diverse set of sequences than IQ-TREE: Hamming distance between reconstructions saturates at 0.2 for the latter, while it steadily increases for the former. Higher diversity can be interpreted as a greater uncertainty concerning the ancestral sequence. However, this must be put in the context of Figure 1: sequences obtained by Bayesian autoregressive reconstruction are more varied but also on average closer to the real ancestor.

**FIG. 2.**
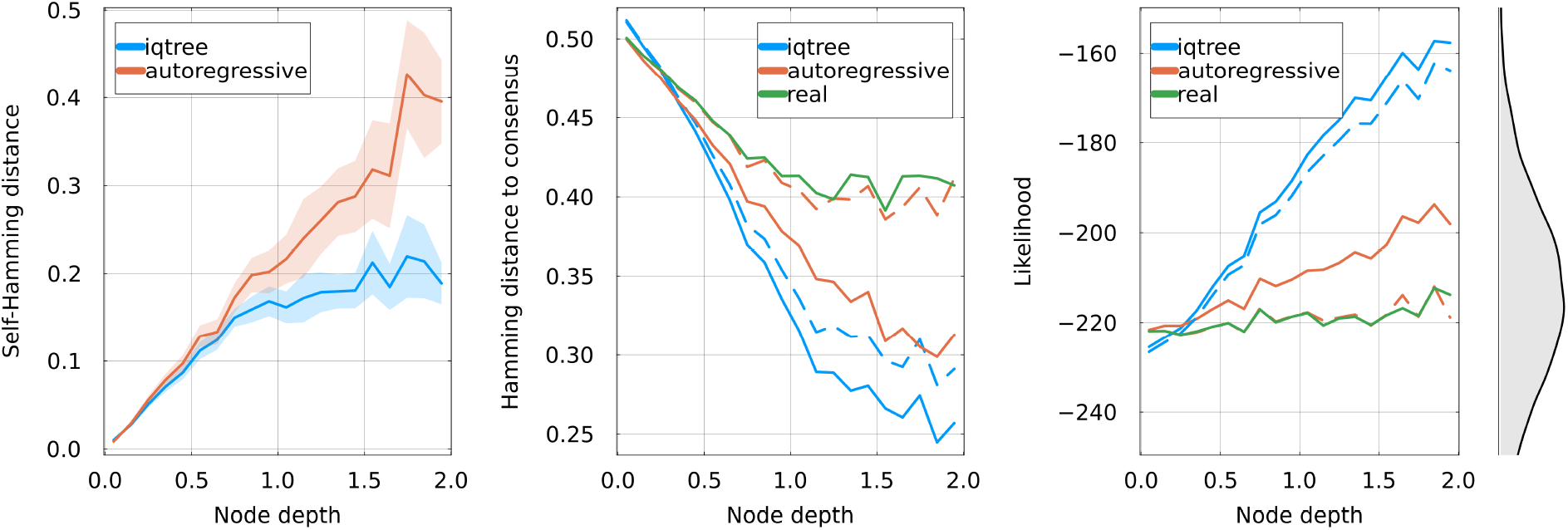
**Left**: for Bayesian reconstruction, average pairwise Hamming distance among sequences reconstructed for each internal node. This quantifies the diversity of sequences obtained using Bayesian reconstruction. **Center**: Hamming distance between reconstructed sequences and the consensus sequence of the alignment. Solid lines represent maximum likelihood reconstruction or the real internal sequences, and dashed lines Bayesian reconstruction. IQ-TREE appears more biased towards the consensus sequence. **Right**: Log-likelihood of reconstructed and real sequences in the autoregressive model, *i*.*e*. using the logarithm of Eq. 2. Maximum likelihood methods (orange and blue solid lines) are biased towards more probable sequences. Bayesian autoregressive reconstruction gives sequences that are at the same likelihood level than the real ancestors. The equilibrium distribution of likelihood of sequences generated by Eq. 2 is shown on the right.

The difference in sequence diversity for the two methods is in part explained by the central panel of Figure 2, which shows the Hamming distance between reconstructed ancestors and the consensus sequence of the multiple sequence alignment at the leaves. It appears there that for deep nodes, IQ-TREE reconstructs sequences that are relatively similar to the consensus, with an average distance between the Bayesian reconstruction and the consensus of about 0.3. Contrasting with that, results of the autoregressive method shows less bias towards the consensus with an average distance of 0.4 for deep nodes, in line with the real ancestors. We also note that maximum likelihood sequences for both method are always closer to the consensus than Bayesian ones, a bias of maximum likelihood already that had already been observed [33].

The bias induced by ignoring the equilibrium distribution of the sequences is also visible in the right panel of Figure 2: it shows the log-likelihood of reconstructed and real ancestral sequences according to the generative model. Note that the log-likelihood here comes from the log-probability of Eq. 2 and can be interpreted as the “quality” of a sequence according to the generative model. It is unrelated to the likelihood computed in Felsenstein’s pruning algorithm. Reconstructions with IQ-TREE increase in likelihood when going deeper in the tree, eventually resulting in “too good” sequences that are very uncharacteristic of the equilibrium generative distribution as can be seen from the histogram on the right. This effect also happens with the maximum likelihood reconstruction of the autoregressive model, although to a lesser extent. The Bayesian autoregressive reconstruction does not suffer from this bias and reconstructs sequences with a log-likelihood that is similar to that of the real ancestors.

#### Potts evolver

We assess the performance of our reconstruction method in the case where the evolver is a Potts model. Potts models are a simple type of generative model and have been used extensively to model protein sequences. They can be use to predict contact in three dimensional structures, effects of mutations, protein-protein interaction partners [10]. They can be sampled to generate novel sequences which are statistically similar to natural ones and often functional [15, 19]. Additionally, it has recently been shown that they can be used to describe the evolution of protein sequences both qualitatively and quantitatively [34].

Potts and autoregressive models both accurately reproduce the statistical properties of protein families. In this sense, they correspond to similar long term generative distributions in the sense of Eq. 5. However, the dynamics of a Potts model are fundamentally different from the ones of usual evolutionary models, including our autoregressive one. Indeed, they are described by a *discrete* time Markov chain, instead of the continuous time used in models based on substitution rate matrices such as in Eq. 1 [19]. For Metropolis steps which we use here, the discrete time corresponds to attempts at mutation which can be either accepted or rejected depending on the effect of the mutation according to the model. These dynamics naturally give rise to different evolutionary timescales for various sequence positions, as well as interesting qualitative behavior such as the entrenchment of mutations [20].

To see how this change in dynamics affects our results, we *(i)* sample a large and varied ensemble of sequences from the Potts model and use it to train an autoregressive model, in a way to guarantee consistent long term distributions between the Potts and autoregressive, and *(ii)* evolve the Potts model along random phylogenies, generating alignments for the leaves and the internal nodes in the same way as above. We then attempt reconstruction of internal nodes using the inferred autoregressive model and IQ-TREE. Figure 3 shows the results of reconstruction, with panels directly comparable to Figure 1. We again see a consistent improvement when using the autoregressive model over IQ-TREE, although of a much smaller amplitude, with an absolute improvement gain in Hamming distance of about 2% for deep internal nodes.

**FIG. 3.**
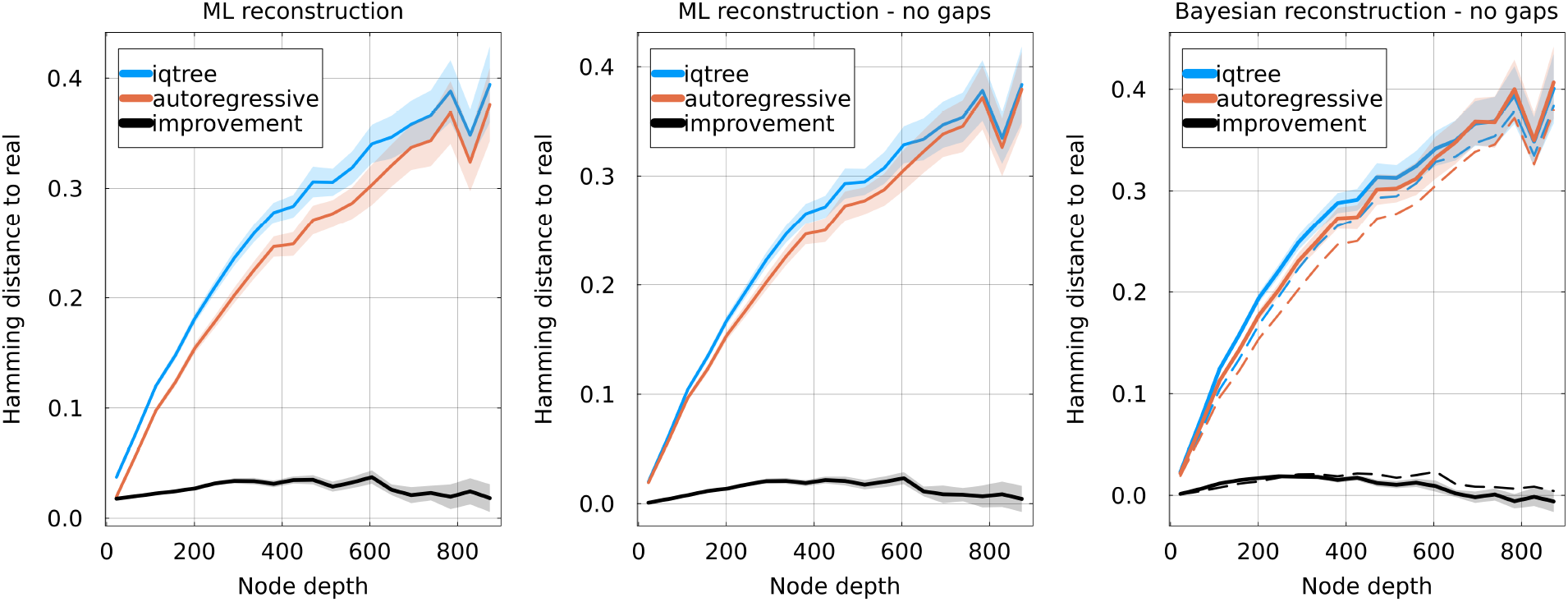
Analogous to Figure 1, but using a Potts model as the evolver. Hamming distance between reconstructed and real sequences as a function of node depth, using IQ-TREE and our autoregressive approach. The difference between the two methods is shown as a black curve. The evolver and reconstruction autoregressive models are learned on the PF00072 family. **Left**: Hamming distance between the full aligned sequences, gaps included, using maximum likelihood reconstruction. **Center**: Hamming distance ignoring gapped positions, using maximum likelihood reconstruction. **Right**: comparison of Bayesian (solid lines) and maximum likelihood (dashed lines) reconstructions, ignoring gaps.

## D. Results on experimental evolution data

We take advantage of recent developments in directed evolution experiments to test our method in a controlled setting. We use the data published in [35]: in this work, authors evolved the antibiotic resistant proteins *β*-lactamase PSE-1 and acetyltransferase AAC6 by submitting them to cycles of mutagenesis and selection for function. Starting from a wild-type protein, they obtained thousands of diverse functional sequences after the directed evolution. An interesting result of this work is that it is possible to recover structural information about the wild-type from the set of evolved sequences.

Here, we use this data as a test setting for ASR: the sequences obtained after directed evolution all derive from a common ancestor, the wild-type, of which we know the amino acid sequence. We can thus reconstruct the wild-type sequence using different ASR methods and compare it to the ground truth. The phylogeny is not known, but given the large population size during the experiment and the relatively low number of selection rounds, it is reasonable to approximate it using a star-tree, *i*.*e*. a tree with a single coalescent event taking place at the root (see Methods). Since the reconstruction task is most interesting when using relatively varied sequences, we decide to use data for the PSE-1 wild-type where 20 cycles of mutagenesis & selection have been performed, resulting in a mean Hamming distance of 12% to the wild-type.

Our ASR procedure is as follows. We randomly pick the amino acid sequences of *M* proteins among the ones evolved from PSE-1 after 20 cycles of mutagenesis & selection, with 3 ≤ *M* ≤ 640. The total number of sequences at round 20 of directed evolution is much larger, making it computationally hard to use all of them. We then construct a star-like phylongeny and place the *M* selected sequences at the leaves, and perform ASR using either IQ-TREE or our autoregressive method which we have trained on an alignment of PSE-1 homologs. We obtain the reconstructed amino acid sequence of the root, which we can then compare to the actual wild-type. As a comparison, and because our approximation of the phylogeny is very simple, we also attempt to reconstruct the root by taking the consensus sequence of the *M* leaves. We repeat this procedure 100 times for each value of *M* for a statistical assessment of the different methods.

The results are shown in Figure 4. The left panel shows the average non-normalized Hamming distance to the wild-type as a function of the number of leaves used *M*. For a low *M*, all methods understandably make a large number of errors, with a mean Hamming distance larger than 10 for *M* = 3. For a higher *M*, IQ-TREE and the autoregressive method stabilize to a fixed number of errors: we find a Hamming distance of ∼ 4.3 for IQ-TREE and ∼ 2.9 for the autoregressive. The consensus curiously reaches a minimum at intermediate *M*, a fact commented in the Supplementary Material, and saturates at a Hamming distance of 6 when considering all sequences of the round 20. The reconstruction errors are overwhelmingly located at six sequence positions. In the central panel, the fraction of mistakes made at these six positions over the 100 repetitions of *M* = 640 leaves is shown for each method. We observe that there are two positions (169 and 193) where IQ-TREE systematically fails at recovering the wild-type state while the autoregressive model’s reconstruction is correct. Interestingly, the corresponding mutations are considered beneficial by the ArDCA model, see Figure S4. Inversely, IQ-TREE recovers the wild-type state more often at position 107. The right panel shows the logo of the set of reconstructed sequences at these 6 positions and for each method.

**FIG. 4.**
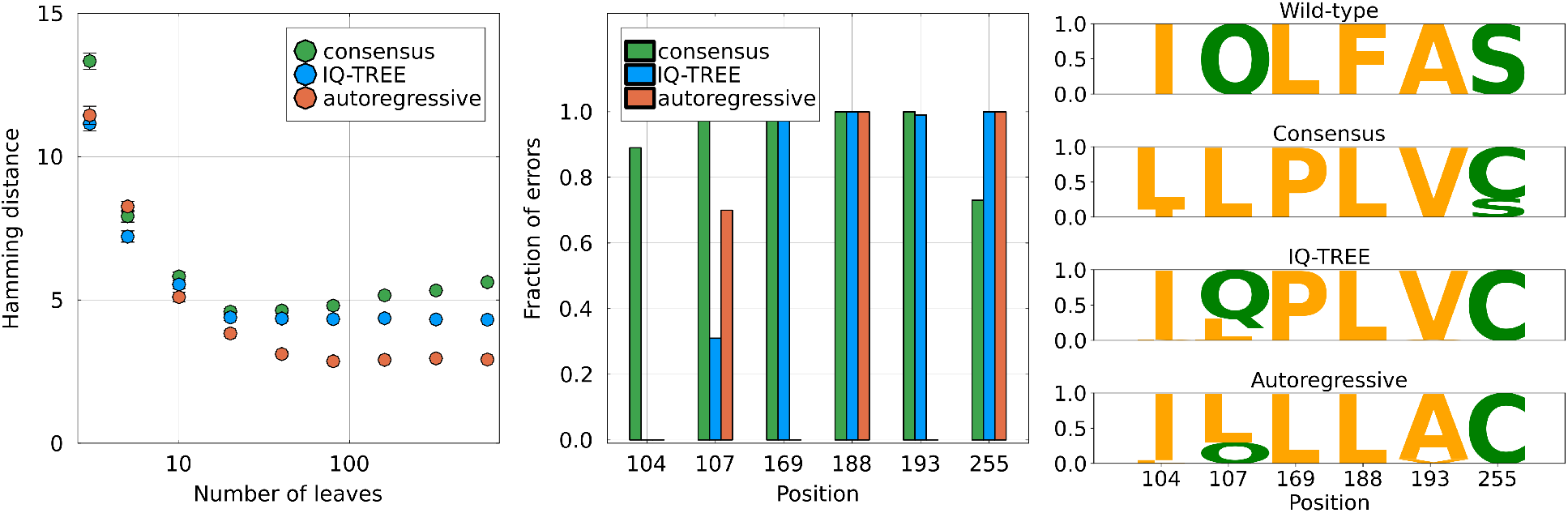
Reconstruction of the wild-type PSE1 sequence used in [35] using sequences from round 20 of the directed evolution. **Left**. Non normalized Hamming distance to the wild-type PSE1 sequence as a function of the number of sequences *M* used for reconstruction. The fact that the consensus method has a local minimum is discussed in the Supplementary Material. For comparison, the average distance between a leaf sequence and the wild-type is 25. The error bars are computed using the standard deviation obtained from the 100 choices of sequences. **Middle**. For the six sequence positions where most of the reconstruction errors are located, fraction of errors of each method out of 100 independent reconstructions using different sets of *M* = 640 leaves. **Right**. Sequence logo of the reconstructed sequence for the three methods, obtained using 100 independent reconstructions with different sets of *M* = 640 leaves. The logo is only shown for the six positions where most errors are located. For example, all three methods fail 100 times at position 147, reconstructing a leucine *L* instead of a phenylalanine *F*.

Overall, we see that the reconstruction of the autoregressive model is more accurate. This gain in accuracy comes from the representation of the functional constraints acting on the PSE-1 protein by the generative model, which are inferred separately using an alignment of homologs. The improvement in reconstruction errors is modest, going from an average Hamming distance of 4.3 to 2.9. However, the gain is intrinsically limited by the data itself: the evolved sequences have an average Hamming distance of about 12% to the ancestor, which is experimentally challenging but remains small compared to the divergence found in the homologs of PSE-1. For instance, the root-to-tip distance estimated by IQ-TREE and the autoregressive model are respectively 0.13 and 0.15, corresponding to the regime of shallow trees when comparing with Figure 1.

## III. DISCUSSION

The reconstruction of ancestral protein sequences has long been a cornerstone of evolutionary biology, helping to elucidate the mechanisms of protein function and evolution over billions of years. The accuracy of ASR has profound implications not only for our understanding of evolution but also for practical applications in synthetic biology and proteins engineering. However, the widely used models in phylogenetics often rely on the assumption of independent sequence evolution at different positions, neglecting epistatic interactions that play a crucial role in determining protein function. This simplification limits their ability to accurately capture the full complexity of evolutionary dynamics.

In this study, we addressed this limitation by developing a novel generative model based on the ArDCA autoregressive framework, which explicitly accounts for epistasis, an essential factor in protein evolution. By incorporating the dependencies between amino acids within sequences, our model offers a more realistic description of protein evolution, capturing the non-independence of mutations over time. A significant contributions of this work is extending the application of generative models to cope with phylogenetic constraints. Our model not only preserves the generative capacity over long-term evolution but it also enables the use of classical phylogenetic techniques such as Felsenstein’s pruning algorithm, which has been a key tool in traditional sequence evolution studies. The ability to integrate a generative context-aware models into these established algorithms represents a substantial advance, allowing for more accurate inference of evolutionary relationships and ancestral states. This, besides the theoretical interest in ASR, is a powerful tool to help us understanding how phylogenetic constraints impact the structure and/or the function of the protein of interest.

Our evaluation of the model using simulated data demonstrated that it outperforms IQ-TREE, a state-of-the-art tool for ASR, in reconstructing ancestral sequences. This improvement highlights the importance of incorporating epistasis into evolutionary models, as ignoring these interactions likely leads to less accurate reconstructions. Furthermore, we validated our approach using experimental data from directed evolution experiments. These data offer a unique opportunity to test the accuracy of ASR methods, and our model achieved more accurate reconstructions of known ancestors compared to IQ-TREE, underscoring the robustness of our approach.

Using the generative nature of our model we can perform Bayesian sampling of sequences at internal nodes that should in principle remain functional despite being distant from any naturally occurring protein. Most ASR studies have used maximum likelihood reconstructions, as Bayesian ones are often found to accumulate to many deleterious mutations and can be non-functional. At the same time, the maximum likelihood solution can be biased and may be unrepresentative of the phenotype of the real ancestor, leading to incorrect biological conclusions [33, 36]. We ourselves observe these biases in our simulations, in the form of a convergence to the consensus sequence and an unnaturally high likelihood according to the generative model. Being able to propose an ensemble of sequences sampled from a generative model at each internal node could thus lead to more robust biological conclusions about ancestral life.

Another feature of our model is its ability to deal with gaps in the multiple sequence alignments in a more principled way compared to other competing algorithms that consider them as missing data. Indeed, in our autoregressive approach gaps are treated as an additional amino acid, and our generative model will produce ancestral sequences that can contain gaps.

The success of our model in both simulations and experimental validation suggests that generative models with autoregressive architectures are powerful tools for studying the dynamics of protein sequence evolution. By capturing the intricacies of epistatic interactions, our model not only improves the accuracy of ancestral sequence reconstruction but also provides new insights into the underlying evolutionary processes. Future work could explore the application of this model to other protein families and further refine the methodology to enhance its applicability in broader phylogenetic contexts.

In conclusion, the integration of epistasis into evolutionary models represents a necessary and timely advancement for the field. Our generative model provides a more nuanced understanding of protein evolution, paving the way for more accurate reconstructions of ancestral sequences and a deeper exploration of the evolutionary dynamics that shape the diversity of life.

## IV. METHODS

### A. ArDCA

The ArDCA model assigns a probability to any sequence of amino acids of length *L* given by

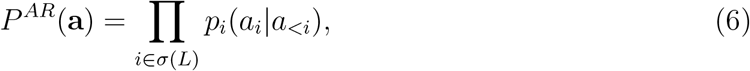

where *σ*(*L*) is a permutation of the *L* first integers and *a*_<*i*_ stands for *a*_1_, …, *a*_*i*−1_. This means that the order in which the conditional probabilities *p*_*i*_ are applied is not necessarily the sequence order. The permutation *σ* is fixed at model inference.

Conditional probabilities *p*_*i*_ are defined as

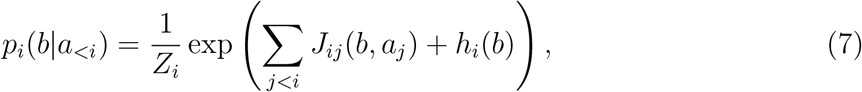

with the *i q*-dimensional vectors *J*_*i*._ and *h*_*i*_ are learned parameters. The model is normally trained using a multiple sequence alignment of homologous proteins, *i*.*e*. a protein family, by finding the parameters *J* and *h* that maximize the likelihood of the sequences. It was shown in [21] that this specific parametrization captures essential features of the variability of members of a protein family.

By definition, homologous proteins share a joint evolutionary history and cannot be considered as statistically independent. To avoid biases, a reweighting is applied to sequences based on their vicinity to other sequences. This scheme has been showed to substantially increase the performance of such models [10].

### B. Branch length inference

With site independent models, optimizing branch lengths of a fixed topology tree to maximize the likelihood of the data is a straightforward task, using for example an algorithm described in [22]. However, this requires an explicit summation over all states at given internal nodes, which we cannot perform using our autoregressive model. The idea we use to reconstruct ancestral sequences, *i*.*e*. inferring the amino acid states of sites one by one in the context of the previous ones, is not applicable here. However, since the dynamics of our autoregressive model is *a priori* quite different from standard models used in IQ-TREE, we prefer to re-infer branch lengths before the ancestral reconstruction.

To work around this issue, we infer branch lengths using a model site-specific frequencies and a unique substitution rate. The site-specific frequencies are chosen so as to match the ones found in a large and long-term sample of our autoregressive model. In this way, we infer branch lengths in a tractable way, using a model that differs from the autoregressive only in the epistatic interactions and not in the conservation profile. Figure S2 shows the good quality of the reconstruction using this technique.

### C. Simulations

A simulation is performed as follows. First, a random tree of *n* = 100 leaves is generated from Yule’s coalescent. We then normalize its height to a fixed value *H* that depends on the evolver model used: for the autoregressive model we use *H* = 2.0, while for the Potts model combined with Metropolis steps, we use *H* = 8 sweeps, *i*.*e. H* = 8*L* Metropolis steps where *L* is the length of the sequences.

A root sequence is sampled from the evolver model’s equilibrium distribution, and evolution is simulated along each branch independently starting from the root. In the case of the autoregressive evolve, the dynamics is the one of Eq. 3. In the case of the Potts model, we use a Markov chain with the Metropolis update rule. In this way, we obtain for each repetition a tree and the alignments for internal and leaf nodes. Results presented in this work are obtained by averaging over *M* = 100 such simulations for each protein family.

### D. Experimental evolution data

To validate the proposed method, we use data from Directed Evolution experiment on Beta-lactamase PSE-1 published in [35]. Beta-lactamase is an enzyme produced by bacteria that provides them with resistance to a certain class of antibiotics known as *betalactam antibiotics*. Its activity relies on the ability to break, through hydrolysis, a certain molecular structure common to all *beta-lactam antibiotics*, the *beta-lactam ring*, inhibiting the antibacterial property of these antibiotics. In [35] the Wild Type (WT) sequence, 266 aminoacids long, has undergone 20 rounds of controlled evolution with an average target mutation rate of approximately 3%-4% per round, and the selected feature is their antibiotic resistance in E.Coli in the presence of ampicillin, slightly above the minimal inhibitory concentration. In these conditions, the percentage of bacterial population that survived selection is approximately 1% (approx. 5 × 10^4^ cells). At round 20, the last one of the experiment, the library of mutated variants has accumulated an average Hamming distance from WT of 12.9% and an average pairwise distance of 19.8%.

A family of 42k homologous sequences is available from PFAM with code PF13354. For this family, an Hidden Markov Model (HMM) of length 214, built on 66 seed sequences, is contextually available. We aligned the experimental sequences to the family HMM according to the following procedure:

1. The WT sequence has been aligned according to the HMM using HMMER.
2. The insertion sites has been removed from the aligned WT sequence.
3. The corresponding insertion sites has been removed from all the other sequences from the experimental library.
4. Gap sites in the aligned WT sequence has been inserted to all the other experimental sequences.

This method ensures that all the aligned positions correspond to the same one for all tested sequences. We remark that in doing this we make the reasonable assumption that all the experimental sequences are aligned to the HMM in same way the WT is aligned to the HMM. It has been noticed in [34] that taking into account the transition possibilities between amino acids allowed by the genetic code is important when describing short term evolutionary dynamics with generative models. In our framework, a natural way to include these is by using the symmetric matrix **H** in the decomposition of Eq. 1. Terms of the **H** matrix do not affect the equilibrium distribution of the model, which thus remains generative, but influences the short term dynamics. Here, we simply counted the number of possibilities to transition from any amino acid to any other based on the genetic code, and we constructed the corresponding **H** matrix. The diagonal matrix remains given by the equilibrium probabilities of amino acids in the context of the sequence, as given by Eq. 4. We found that this substantially improves the results of the autoregressive reconstruction for the experimental evolution data.

### E. Reconstruction with IQ-TREE

We run IQ-TREE using the `-asr` flag to generate states at internal nodes of the tree. By default, IQ-TREE reconstructs the maximum likelihood sequence at internal nodes. It also generates a “state” file containing the posterior probabilities of amino acids at each internal node, from which we sample to obtain the Bayesian reconstruction.

On simulated data, we ran IQ-TREE using the model finder routine to select the evolutionary model [37]. The model most frequently found was based on the PMB matrix [38], with different options for rates, *e*.*g*. `+G4, +I+G4 or +R4.` On the directed evolution data, the two most frequently found models were JTT [3] and the between patient HIV model of [39]. Since the latter is clearly unrelated to the protein that is considered here, we used the `JTT+G4` model for reconstruction.

### F. Code & data availability

The code used in this work is accessible at the following links:

- the implementation of the reconstruction algorithm described here is available at https://github.com/PierreBarrat/AncestralSequenceReconstruction.jl
- the code used in simulations and data analysis is available at https://github.com/PierreBarrat/AutoRegressiveASR.

## Acknowledgments

We thank Juan Rodriguez-Rivas for useful discussions.

## Appendix A Autoregressive evolution model

### 1. Simplified expression for a homogeneous H

For each site *i*, the main difference between our model and a traiditional GTR is that the equilibrium frequencies of the Markov chain are computed using the context at the previous sites 1, …, *i* − 1. Considering Eq. 1 and Eq. 4, this means that the diagonal matrix is determined using the generative model. On the other hand, the symetric matrix **H** can be given any value without changing the long term generative properties of the dynamical model, *i*.*e*. Eq. 5. Here, we show that if the transitions defined by **H** are uniform, *i*.*e. H*_*ab*_ = *µ* for any *a* ≠ *b*, the propagator takes a simplified form:

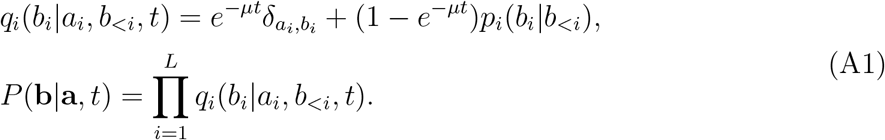

The interpretation of the site propagator *q*_*i*_(*b*_*i*_|*a*_*i*_, *b*_<*i*_, *t*) is straightforward: if no mutation occurs with probability *e*^−*µt*^, site *i* remains in its original state *a*_*i*_; otherwise, with probability (1 − *e*^−*µt*^), it is resampled using the equilibrium probability given by the generative model and the context of the sequence *p*_*i*_(*b*_*i*_|*b*_<*i*_). Note that the assumption of a scalar matrix is reasonable if one wishes to ignore the different transition rates between amino-acids.

To lighten notation, we drop the explicit dependence on the position *i* and the sequence context *b*_<*i*_ by defining *p*_*b*_ = *p*_*i*_(*b*_*i*_|*b*_<*i*_). We will then compute the *n* eigenvectors and eigenvalues of **Q**, where *n* = 21 for the amino acids and gap symbol. First, note that for the continuous time Markov chain to be well defined, we need the rows of **Q** to sum to 0. We thus have the following expression for the elements of **Q**:

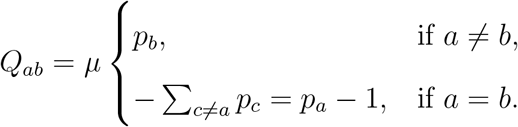

This means that the vector 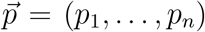 is the left-eigenvector of **Q** associated to eigenvalue 0:

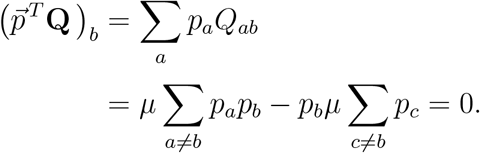

Since the dynamics of the Markov chain are given by 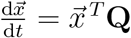, this implies that 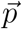 is an equilibrium state of the chain, as expected.

We now prove that the other *n* − 1 eigenvalues are equal to −*µ*. Consider a vector 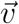 orthogonal to 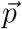 The following calculation shows that 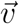 is a right-eigenvector of **Q** with eigenvalue −*µ*:

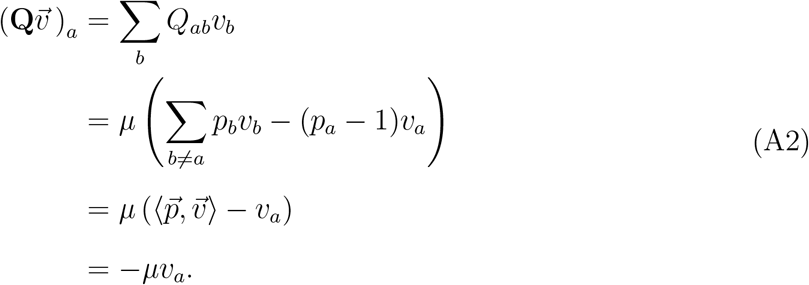

The space of vectors orthogonal to 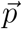 is of dimension *n* − 1, therefore eigenvalue −*µ* is degenerate and the corresponding eigenvectors span a space of dimension *n* − 1. We call 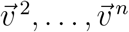 an orthogonal basis of this space. As the rows of **Q** sum to 0, the remaining eigenvector is 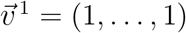 and is associated to eigenvalue 0. Similarly, we can also show that there exist *n*− 1 left-eigenvectors associated to the eigenvalue −*µ*, which note 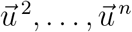, defined by the relation 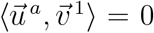. The first left-eigenvector is of course 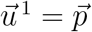.

With the right choice of the basis 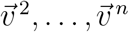 and 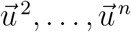, we have the relation 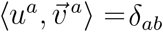 This means that we can write **Q** as

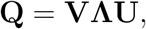

where **V** has the right eigenvectors as columns, **U** the left eigenvectors as rows, and

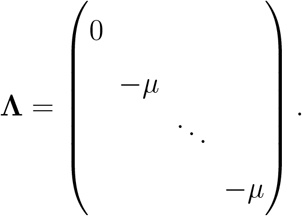

We now compute the transition probability *q*(*b*|*a, t*) by computing the exponential of **Q**:

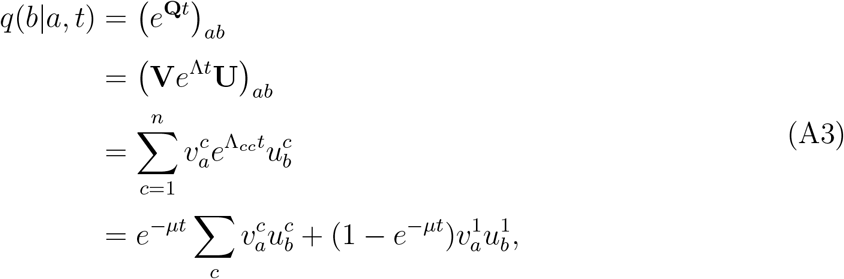

where we have used the fact that Λ_*cc*_ = −*µ* for *c* > 1 and 0 for *c* = 1. Using the fact that **V** and **U** are each other’s inverse, we have that 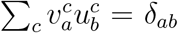. Furthermore, 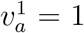 and 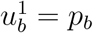 We thus obtain the desired result:

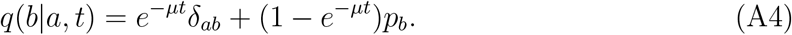

### 2. Irreversibility of the autoregressive evolution model

For each position *i* and given a sequence context, the autoregressive model has the same structure as classical sequence evolution models. In particular, it is time reversible: given any two amino acid states *a*_*i*_ and *b*_*i*_, there is no objective way of determining whether *a*_*i*_ evolved in to *b*_*i*_ or the reverse. However, the propagator for full sequences Eq. 3 is *time irreversible*. We demonstrate this using a simple example.

For a stochastic model, time reversibility is equivalent to respecting *detailed balance*: for any two sequences **a** and **b** and any time *t*, one should have

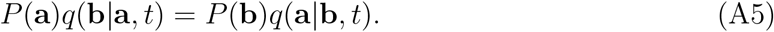

We set out to design a simple autoregressive model that does not fulfill detail balance, showing that time reversibility is in general not fulfilled. Consider a sequence of length *L* = 2 with binary states *a*_1_, *a*_2_ ∈ {0, 1}. Assume that the “fitness landscape” of this sequence is such that sequences (0, 0) and (1, 1) are functional, while (0, 1) and (1, 0) are not. A corresponding sequence alignment would show the following distribution of sequences:

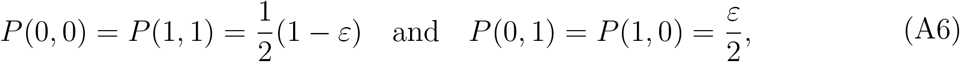

with *ε* ≪ 1. A well trained autoregressive model would consequently have the following properties:

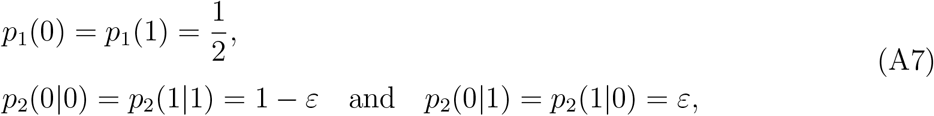

which is the same distribution as in Eq. SA6.

We now consider the two sequences **a** = (1, 1) and **b** = (0, 1) to show that the model does not respect detailled balance. Computing the two sides of Eq. SA5 in this configuration, we find

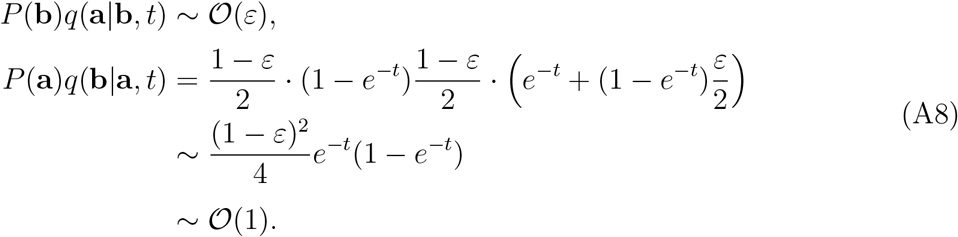

The first expression is of order *ε* since the probability that the process started in **b** is *P*(**b**) = *ε/*2; even if *q*(**a**|**b**, *t*) is a likely transition, the scenario **b** → **a** remains unlikely. For the second expression however, things are different. The right hand side is composed successively of *P*(**a**) = (1 − *ε*)*/*2, of *q*_1_(0|1, *t*) = (1 − *e*^−*t*^)(1 − *ε*)*/*2, and of *q*_2_(1|1, *b*_<2_ = (0), *t*) = *e*^−*t*^ + (1 − *e*^−*t*^)*ε/*2. This shows that the scenario **a** → **b** is more likely, and we can assume that **a** is the ancestor of **b**.

Two things can now be discussed. First, even though the model does not satisfy detailed balance, it is straightforward to show that it satisfies *global balance* and thus that it has a well defined equilibrium distribution. For instance, in the toy model above, the long term distribution of the propagator *q* can be shown to be equal to the one in Eq. SA6. Secondly, the cause of irreversibility here is not epistasis in itself, but rather the structure of the autoregressive probability distribution. In our toy example, the model eagerly mutates the first residue without considering the second one, allowing a likely **a** → **b** transition. In fact, it is perfectly possible to design dynamical epistatic models that are time reversible [19]. Furthermore, irreversible models of evolution are not the most common but are still part of standard phylogenetic toolkits [29].

## Appendix B Directed evolution data

### 1. Minimum reconstruction error of the consensus

In the left panel of Figure 4, the Hamming distance of the consensus of *M* sequences to the wild-type sequence shows a minimum for an intermediate value of *M*. This is at first counter-intuitive, and we present here a minimalistic example to illustrate this phenomenon.

We consider the simplified case of a star-like tree with *M* leaves. The root sequence is **y** = (*y*_1_, …, *y*_*L*_) with *y*_*i*_ = 0, and the sequence of leaf *m* is **x**^*m*^ = (*x*_1_, …, *x*_*L*_) with *x*_*i*_ ∈ {0, 1}. We now assume that during evolution, the first site has mutated with probability *p* = 1*/*2 + *ε* while the remaining *L* − 1 have mutated with probability *q* = *ε*, with *ε* < 1*/*2. For site 1 (resp. for the remaining sites), the probability that the consensus of the leaves differs from the root is equal to the probability that a binomial variable of parameters (*p, M*) (resp. (*q, M*)) takes a value larger than *M/*2. We call *α*_*p*_(*M*) (resp. *α*_*q*_(*M*)) this probability. It is immediate that the average Hamming distance *H*(*M*) between the consensus and the root if there are *M* leaves is

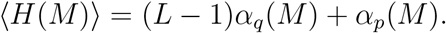

The limits of *α* for large and small *M* are easily obtained:

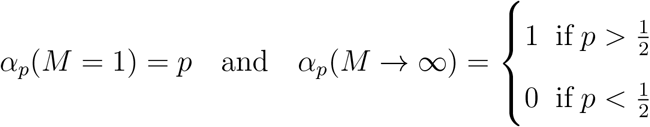

We therefore obtain the limits of ⟨*H*(*M*)⟩:

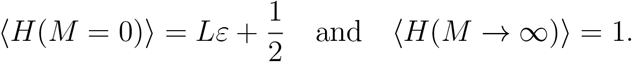

It is interesting to note that depending on *ε* and *L*, the consensus of *M* = 1 sequence can be on average closer to the ancestral sequence than the consensus of a very large number of sequences. Unfortunately, there are no convenient analytical expressions for *α*_*q*_ and *α*_*p*_ for intermediate *M*. In Figure S1, we show the numerical values of these two quantities for a specific choice of *ε*: *α*_*p*_ increases monotonically since *p* > 1*/*2, while *α*_*q*_ decreases since *q* < 1*/*2. The average Hamming distance ⟨*H*(*M*)⟩ is then observed to have a minimum for an intermediate *M*.

**Figure S 1.**
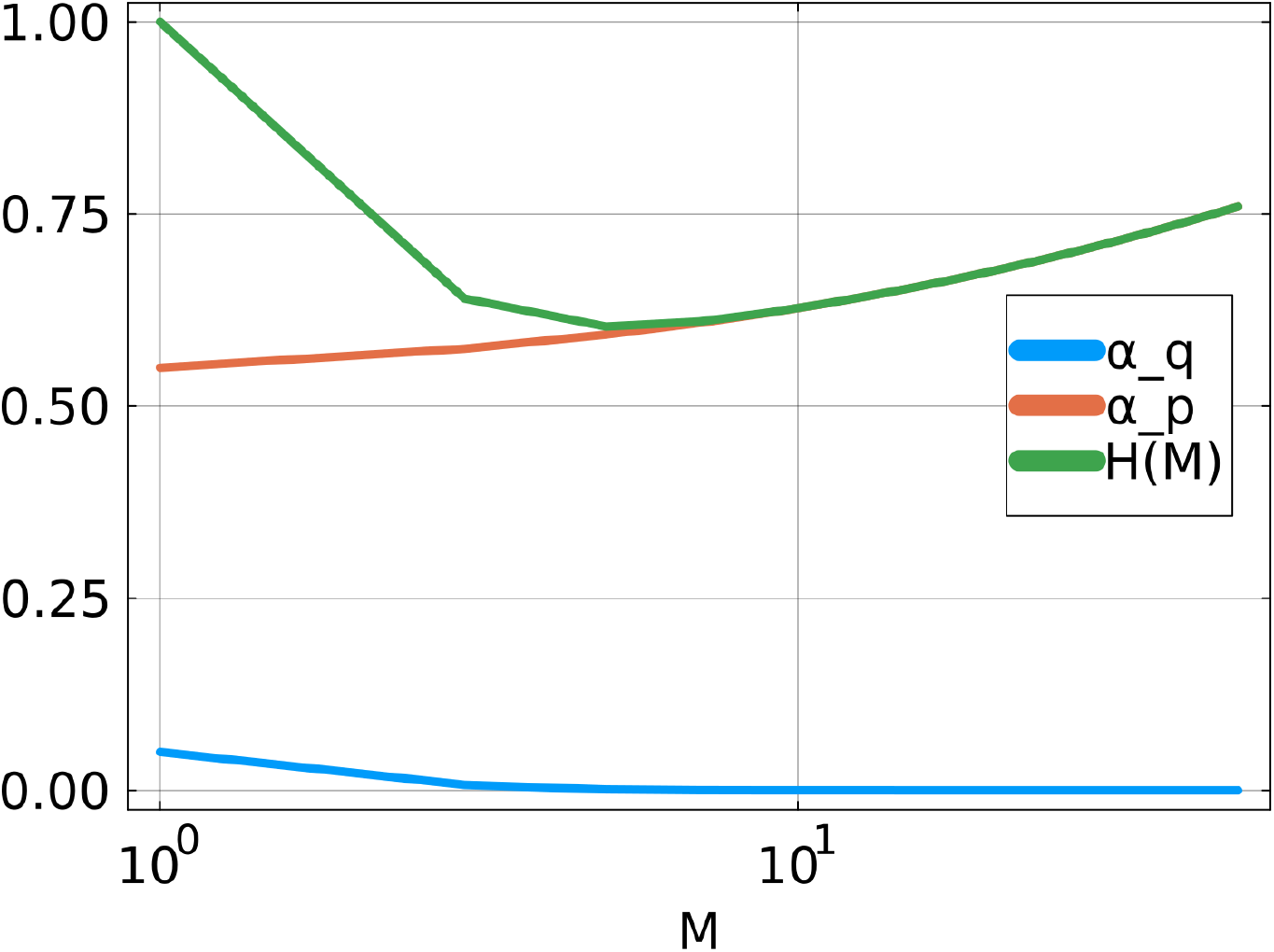
Quantities *α*_*p*_, *α*_*q*_ and ⟨*H*⟩ as a function of the number of leaves *M* (odd values only). *α*_*p*_ is increasing from 1*/*2 + *ε* to 1 while *α*_*q*_ is decreasing from *ε* to 0. The average Hamming distance (*L* − 1)*α*_*q*_(*M*) + *α*_*p*_(*M*) reaches a minimum for an intermediate number of leaves. Values of parameters: *ε* = 0.05, *L* = 10.

## Appendix C: Supplementary figures

**Figure S 2.**
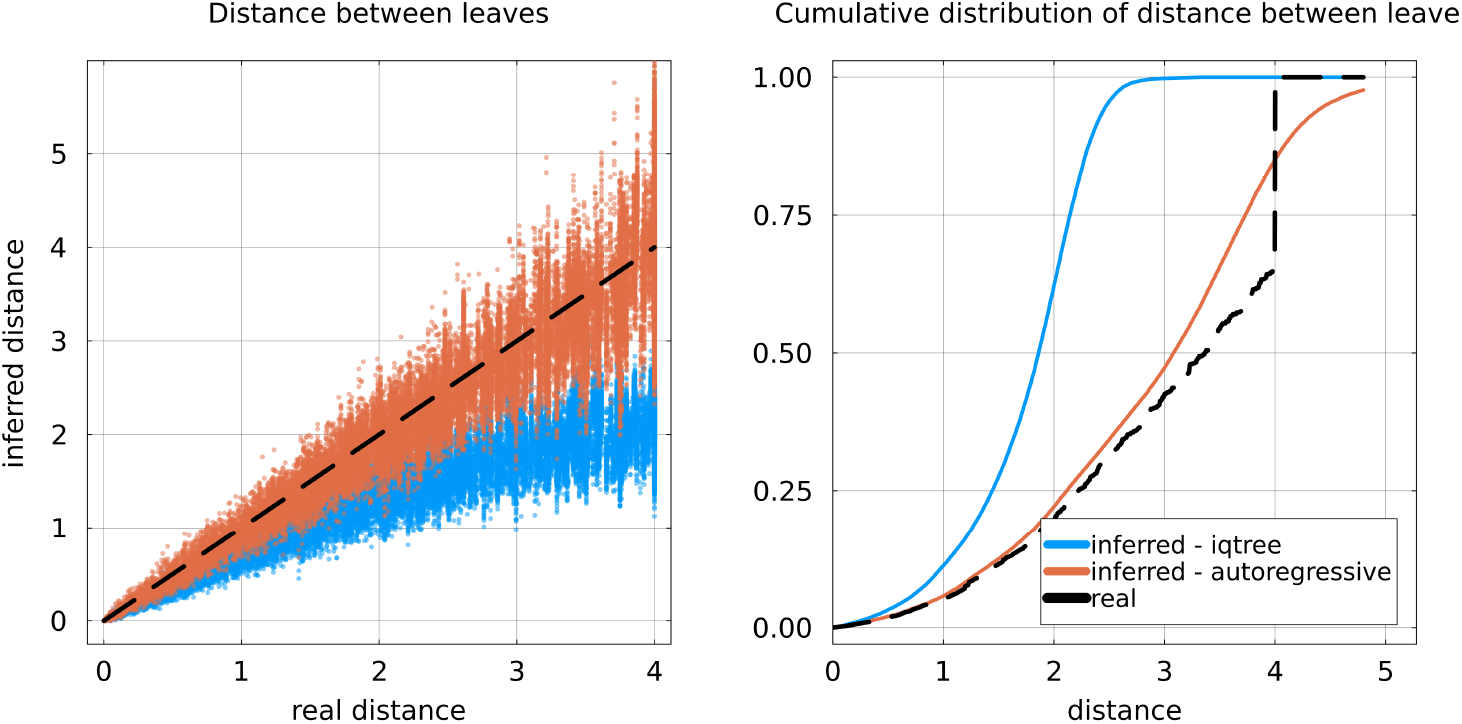
Quality of branch length inference using data simulated with the autoregressive evolver and a tree with fixed topology. Two techniques are compared: IQ-TREE and the profile model corresponding to the autoregressive evolver. **Left**: inferred distance vs distance in the real trees for every pair of leaves. **Right**: Cumulative distribution of pairwise distance along the tree between leaves for the two inference methods and for the real tree. The discontinuity in the curve for the real tree is caused by the ultrametricity and fixed total height of the generated trees.

**Figure S 3.**
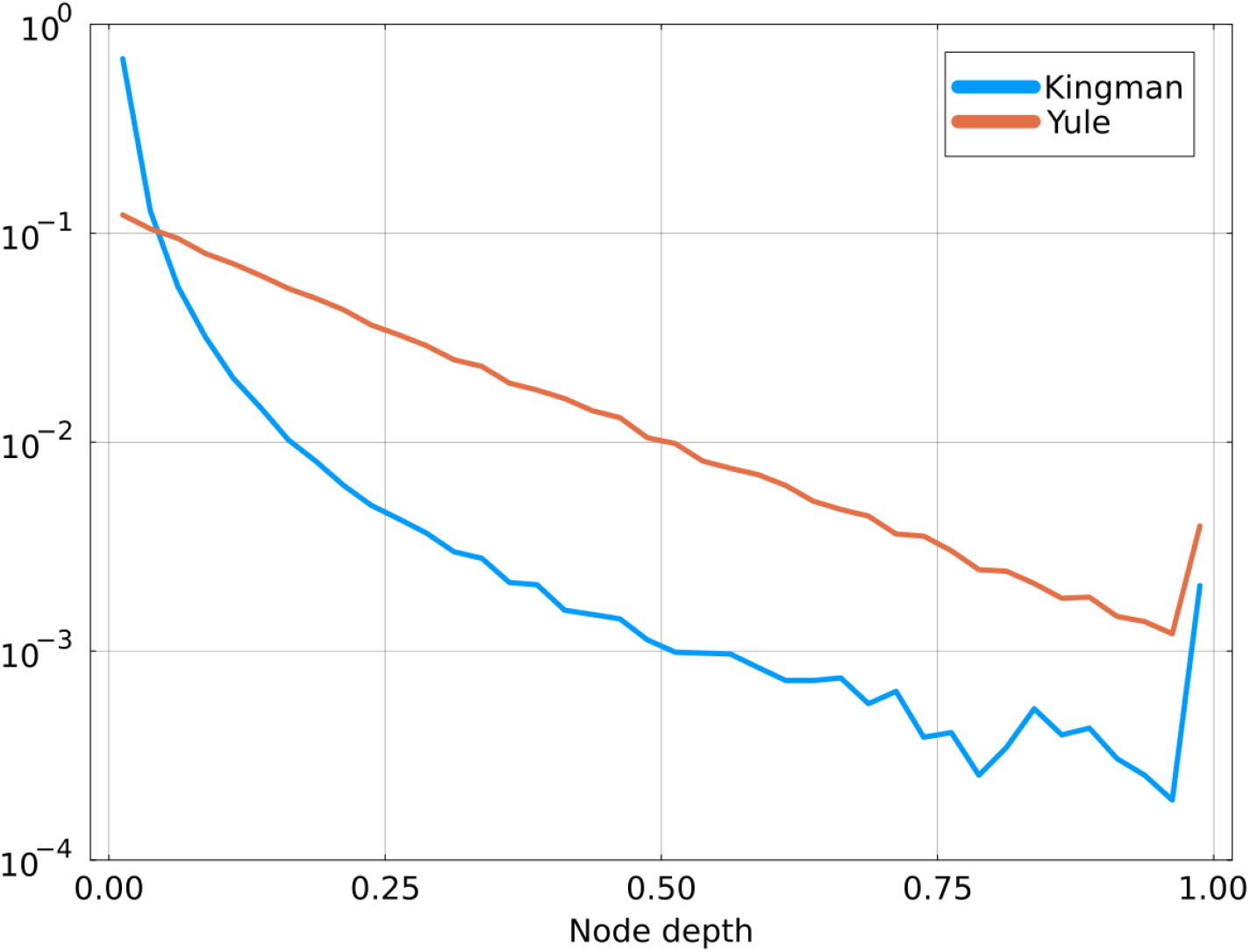
Distribution of node depth for trees coming from the Kingman and Yule coalescents. Node depth is defined as the distance from a node to the closest leaf. Data is obtained by sampling several trees from each coalescent. Heights of trees are normalized to one. The Kingman process concentrates most of the nodes in close vicinity to the leaves, while the Yule process spreads them more evenly.

**Figure S 4.**
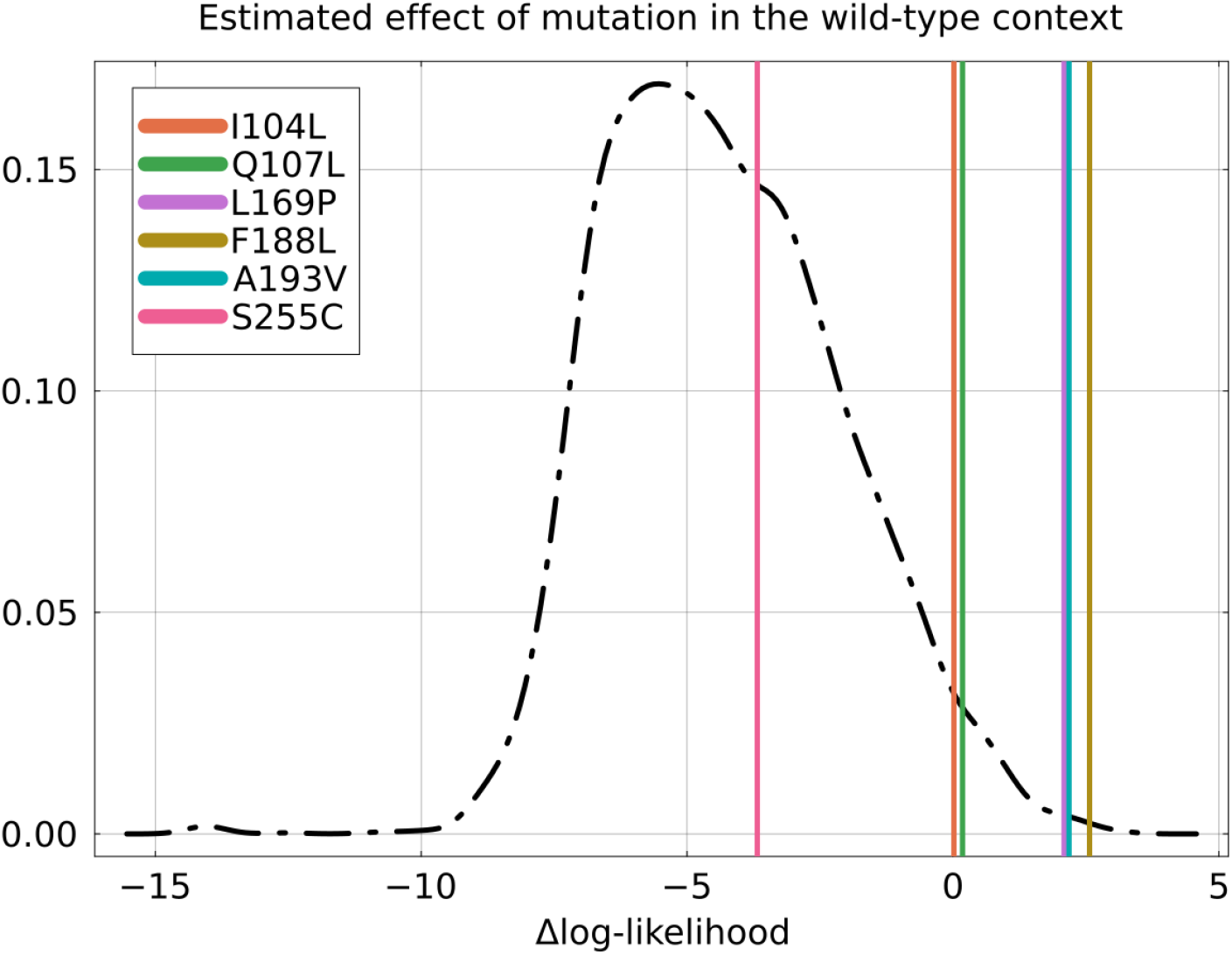
Distribution of estimated effect of single mutations by ArDCA in the PSE1 sequence (black curve). The effect of a mutations is estimated by computing the difference in log-likelihood between the mutant sequence and the wild-type: negative values are detrimental and 0 represents a neutral mutation. As expected, most mutations are estimated to be detrimental but mutations found in the consensus of round 20 are mostly beneficial or neutral. The six reconstruction errors in Figure 4 are displayed as vertical bars. The two positions 169 and 193 where ArDCA outperforms IQ-TREE correspond to beneficial mutations.

